# FactorNet: a deep learning framework for predicting cell type specific transcription factor binding from nucleotide-resolution sequential data

**DOI:** 10.1101/151274

**Authors:** Daniel Quang, Xiaohui Xie

## Abstract

Due to the large numbers of transcription factors (TFs) and cell types, querying binding profiles of all TF/cell type pairs is not experimentally feasible, owing to constraints in time and resources. To address this issue, we developed a convolutional-recurrent neural network model, called FactorNet, to computationally impute the missing binding data. FactorNet trains on binding data from reference cell types to make accurate predictions on testing cell types by leveraging a variety of features, including genomic sequences, genome annotations, gene expression, and single-nucleotide resolution sequential signals, such as DNase I cleavage. To the best of our knowledge, this is the first deep learning method to study the rules governing TF binding at such a fine resolution. With FactorNet, a researcher can perform a single sequencing assay, such as DNase-seq, on a cell type and computationally impute dozens of TF binding profiles. This is an integral step for reconstructing the complex networks underlying gene regulation. While neural networks can be computationally expensive to train, we introduce several novel strategies to significantly reduce the overhead. By visualizing the neural network models, we can interpret how the model predicts binding which in turn reveals additional insights into regulatory grammar. We also investigate the variables that affect cross-cell type predictive performance to explain why the model performs better on some TF/cell types than others, and offer insights to improve upon this field. Our method ranked among the top four teams in the ENCODE-DREAM *in vivo* Transcription Factor Binding Site Prediction Challenge.

## Introduction

High-throughput sequencing has led to a diverse set of methods to interrogate the epigenetic landscape for the purpose of discovering tissue and cell type-specific putative functional elements. Such information provides valuable insights for a number of biological fields, including synthetic biology and translational medicine. Among these methods are ChIP-seq, which applies a large-scale chromatin immunoprecipitation assay that maps *in vivo* transcription factor (TF) binding sites or histone modifications genome-wide (Johnson et al., 2007), and DNase-seq, which identifies genome-wide locations of open chromatin, or “hotspots”, by sequencing genomic regions sensitive to DNase I cleavage (Crawford, G. et al., 2006; John et al., 2013). At deep sequencing depth, DNase-seq can identify TF binding sites, which manifest as dips, or “footprints”, in the digital DNase I cleavage signal (Hesselberth et al., 2009; Boyle et al., 2011; Neph, S. et al., 2012). Other studies have shown that cell type-specific functional elements can display unique patterns of motif densities and epigenetic signals (Quang et al., 2015b). Computational methods can integrate these diverse datasets to elucidate the complex and non-linear combinations of epigenetic markers and raw sequence contexts that underlie functional elements such as enhancers, promoters, and insulators. Some algorithms accomplish this by dividing the entire genome systematically into segments, and then assigning the resulting genome segments into “chromatin states” by applying machine learning methods such as Hidden Markov Models, Dynamic Bayesian Networks, or Self-Organizing Maps (Ernst and Kellis, 2012; Hoffman et al., 2012; Mortazavi et al., 2013).

The Encyclopedia of DNA Elements (ENCODE) (ENCODE Project Consortium, 2012) and NIH Roadmap Epigenomics (Roadmap Epigenomics Consortium et al., 2015) projects have generated a large number of ChIP-seq and DNase-seq datasets for dozens of different cell and tissue types. Owing to several constraints, including cost, time or sample material availability, these projects are far from completely mapping every mark and sample combination. This disparity is especially large for TF binding profiles because ENCODE has profiled over 600 human biosamples and over 200 TFs, translating to over 120,000 possible pairs of biosamples and TFs, but as of the writing of this article only about 8,000 TF binding profiles are available. Due to the strong correlations between epigenetic markers, computational methods have been proposed to impute the missing datasets. One such imputation method is ChromImpute (Ernst and Kellis, 2015), which applies ensembles of regression trees to impute missing chromatin marks. With the exception of CTCF, ChromImpute does not impute TF binding. Moreover, ChromImpute does not take sequence context into account, which can be useful for predicting the binding sites of TFs like CTCF that are known to have a strong binding motif.

Computational methods designed to predict TF binding include PIQ (Sherwood et al., 2014), Centipede (Pique-Regi et al., 2011), and msCentipede (Raj et al., 2015). These methods require a collection of motifs and DNase-seq data to predict TF binding sites in a single tissue or cell type. While such an approach can be convenient because the DNase-seq signal for the cell type considered is the only mandatory experimental data, it has several drawbacks. These models are trained in an unsupervised fashion using algorithms such as expectation maximization (EM). From our experience, EM-based algorithms can be very computationally inefficient. To compensate for this issue, PIQ, Centipede, and msCentipede limit training and evaluation to motif matches, which represent a small and unrepresentative fraction of the whole genome. Furthermore, the manual assignment of a motif for each TF is a strong assumption that completely ignores any additional sequence contexts such as co-binding, indirect binding, and non-canonical motifs. This can be especially problematic for TFs like REST, which is known to have eight non-canonical binding motifs (Quang and Xie, 2014).

More recently, deep neural network (DNN) methods have gained significant traction in the bioinformatics community. DNNs are useful for biological applications because they can efficiently dentify complex non-linear patterns from large amounts of feature-rich data. They have been successfully applied to predicting splicing patterns (Leung et al., 2014), predicting variant deleteriousness (Quang et al., 2015a), and gene expression inference (Chen et al., 2016). The convolutional neural network (CNN), a variant of the DNN, has been useful for genomics because it can process raw DNA sequences and the kernels are analogues to position weight matrices (PWMs), which are popular models for describing the sequence-specific binding pattern of TFs. Examples of genomic application of CNNs include DanQ(Quang and Xie, 2016), DeepSEA (Zhou and Troyanskaya, 2015), Basset (Kelley et al., 2016), DeepBind (Alipanahi et al., 2015), and DeeperBind (Hassanzadeh and Wang, 2016). These methods accept raw DNA sequence inputs and are trained in a supervised fashion to discriminate between the presence and absence of epigenetic markers, including TF binding, open chromatin, and histone modifications. Consequently, these algorithms are not suited to the task of predicting epigenetic markers across cell types. Instead, they are typically designed for other tasks such as motif discovery or functional variant annotation. Both DanQ and DeeperBind, unlike the other three CNN methods, also use a recurrent neural network (RNN), another type of DNN, to form a CNN-RNN hybrid architecture that can outperform pure convolutional models. RNNs have been useful in other machine learning applications involving sequential data, including phoneme classification (Graves and Schmidhuber, 2005), speech recognition (Graves et al., 2013), machine translation (Sundermeyer et al., 2014), and human action recognition (Zhu et al., 2016). More recently, CNNs and RNNs have been used for predicting single-cell DNA methylation states (Angermueller et al., 2017).

To predict cell type-specific TF binding, we developed FactorNet, which combines elements of the aforementioned algorithms. FactorNet trains a DNN on data from one or more reference cell types for which the TF or TFs of interest have been profiled, and this model can then predict binding in other cell types. The FactorNet model builds upon the DanQ CNN-RNN hybrid architecture by including additional real-valued coordinated-based signals such as DNase-seq signals as features. FactorNet is similar to a recently developed method called DeepCpG, which integrates sequence context and neighboring methylation rates to predict single-cell DNA methylation states using a CNN and a bidirectional RNN (Angermueller et al., 2017). We also extended the DanQ network into a “Siamese” architecture that accounts for reverse complements (Figure 1). This Siamese architecture applies identical networks to both strands to ensure that both the forward and reverse complement sequences return the same outputs, essentially halving the total amount of training data, ultimately improving training efficiency and predictive accuracy. Both networks share the same weights. Siamese networks are popular among tasks that involve finding similarity or a relationship between two comparable objects. Two examples are signature verification (Bromley et al., 1993) and assessing sentence similarity (Mueller and Thyagarajan, 2016). Another recent method, TFImpute (Qin and Feng, 2017), shares many similarities with FactorNet. Like FactorNet, TFImpute is intended to impute missing TF binding datasets, and it uses a CNN-RNN architecture, and a weight-sharing strategy to handle reverse complements. TFImpute is a sequence-only method and therefore more comparable to DeepSEA, DeepBind, and Basset. Unlike FactorNet, TFImpute does not directly accept cell type-specific data like DNase-seq as model inputs.

**Figure 1.**
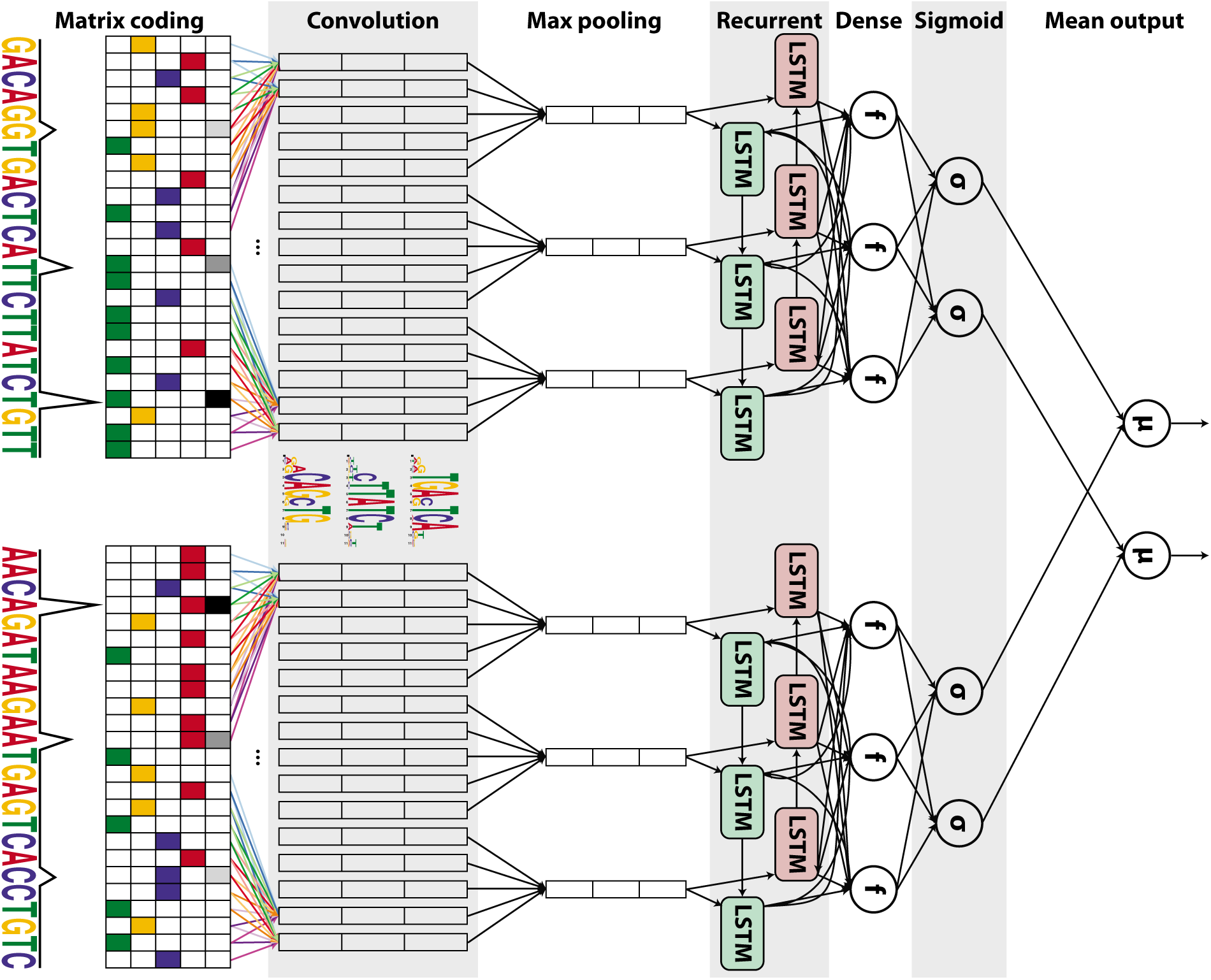
Simplified diagram of the FactorNet model. An input DNA sequence (top) is first one hot encoded into a 4-row bit matrix. Real-valued single-nucleotide signal values are concatenated as extra rows to this matrix. A rectifier activation convolution layer transforms the input matrix into an output matrix with a row for each convolution kernel and a column for each position in the input (minus the width of the kernel). Each kernel is effectively a sequence motif. Max pooling downsamples the output matrix along the spatial axis, preserving the number of channels. The subsequent recurrent layer contains long short term memory (LSTM) units connected end-to-end in both directions to capture spatial dependencies between motifs. Recurrent outputs are densely connected to a layer of rectified linear units. The activations are likewise densely connected to a sigmoid layer that nonlinear transformation to yield a vector of probability predictions of the TF binding calls. An identical network, sharing the same weights, is also applied to the reverse complement of the sequence (bottom). Finally, respective predictions from the forward and reverse complement sequences are averaged together, and these averaged predictions are compared via a loss function to the true target vector. Although not pictured, we also include a sequence distributed dense layer between the convolution and max pooling layer to capture higher order motifs.

We submitted the FactorNet model to the ENCODE-DREAM *in vivo* Transcription Factor Binding Site Prediction Challenge (https://www.synapse.org/ENCODE), where it placed among the top four ranked teams. All results discussed in this paper are derived from data in the Challenge. The Challenge delivers a crowdsourcing approach to figure out the optimal strategies for solving the problem of TF binding prediction.

## Results

### Predictive performance varies across transcription factors

Table 1 shows a partial summary of FactorNet cross-cell type predictive performances on a variety of cell type and TF combinations as of the conclusion of the ENCODE-DREAM Challenge. Final rankings in the Challenge are based on performances over 13 TF/cell type pairs. A score combining several primary performance measures is computed for each pair. In addition to the 13 TF/cell type pairs for final rankings, there are 28 TF/cell type “leaderboard” pairs. Competitors can compare performances and receive live updating of their scores for the leaderboard TF/cell type pairs. Scores for the 13 final ranking TF/cell type pairs were not available until the conclusion of the challenge. Our model achieved first place on six of the 13 TF/cell type final ranking pairs, the most of any team.

**Table 1.** Partial summary of FactorNet cross-cell type predictive performances on the ENCODE-DREAM Challenge data. Each final ranking TF/cell type pair is demarcated with a *. For each final ranking TF/cell type pair, we provide, in parentheses, performance scores based on the evaluation pair’s original ChIP-seq fold change signal.

| Factor | Cell type | auROC | auPR | Recall at 50% FDR |
| --- | --- | --- | --- | --- |
| CTCF* | iPSC | 0.9966 (0.9998) | 0.8608 (0.9794) | 0.9142 (0.9941) |
| CTCF | GM12878 | 0.9968 | 0.8451 | 0.8777 |
| CTCF* | PC-3 | 0.9862 (0.9942) | 0.7827 (0.8893) | 0.7948 (0.9272) |
| ZNF143 | K562 | 0.9884 | 0.6957 | 0.7303 |
| MAX | MCF-7 | 0.9956 | 0.6624 | 0.8290 |
| MAX* | liver | 0.9882 (0.9732) | 0.4222 (0.6045) | 0.3706 (0.6253) |
| EGR1 | K562 | 0.9937 | 0.6522 | 0.7312 |
| EGR1* | liver | 0.9856 (0.9741) | 0.3172 (0.5306) | 0.2164 (0.5257) |
| HNF4A* | liver | 0.9785 (0.9956) | 0.6188 (0.8781) | 0.6467 (0.9291) |
| MAFK | K562 | 0.9946 | 0.6176 | 0.6710 |
| MAFK | MCF-7 | 0.9906 | 0.5241 | 0.5391 |
| GABPA | K562 | 0.9957 | 0.6125 | 0.6299 |
| GABPA* | liver | 0.9860 (0.9581) | 0.4416 (0.5197) | 0.3550 (0.5202) |
| YY1 | K562 | 0.9945 | 0.6078 | 0.7393 |
| TAF1 | HepG2 | 0.9930 | 0.5956 | 0.6961 |
| TAF1* | liver | 0.9892 (0.9657) | 0.4283 (0.4795) | 0.4039 (0.4766) |
| E2F6 | K562 | 0.9885 | 0.5619 | 0.6455 |
| REST | K562 | 0.9958 | 0.5239 | 0.5748 |
| REST* | liver | 0.9800 (0.9692) | 0.4122 (0.5596) | 0.4065 (0.5945) |
| FOXA1* | liver | 0.9862 (0.9813) | 0.4922 (0.6546) | 0.4889 (0.6728) |
| FOXA1 | MCF-7 | 0.9638 | 0.4487 | 0.4613 |
| JUND | H1-hESC | 0.9948 | 0.4098 | 0.3141 |
| JUND* | liver | 0.9765 (0.9825) | 0.2649 (0.6921) | 0.1719 (0.7223) |
| TCF12 | K562 | 0.9801 | 0.3901 | 0.3487 |
| STAT3 | GM12878 | 0.9975 | 0.3774 | 0.3074 |
| NANOG* | iPSC | 0.9885 (0.9876) | 0.3539 (0.6421) | 0.3118 (0.6680) |
| CREB1 | MCF-7 | 0.9281 | 0.3105 | 0.2990 |
| E2F1* | K562 | 0.9574 (0.9888) | 0.2406 (0.6428) | 0.0000 (0.6573) |
| FOXA2* | liver | 0.9773 (0.9932) | 0.2172 (0.7920) | 0.0231 (0.8278) |

FactorNet typically achieves auROC scores above 97% for most of the TF/cell type pairs, reaching as low as 92.8% for CREB1/MCF-7. auPR scores, in contrast, display a wider range of values, reaching as low as 21.7% for FOXA1/liver and 87.8% for CTCF/iPSC. For some TFs, such as CTCF and ZNF143, the predictions are already accurate enough to be considered useful. Much of the variation in auPR scores can be attributed to noise in the ChIP-seq signal used to generate the evaluation labels, which we demonstrate by building classifiers based on taking the mean in a 200 bp window of the ChIP-seq fold change signal with respect to input control. Peak calls are derived from the SPP algorithm (Kharchenko et al., 2008), which uses the fold-change signal and peak shape to score and rank peaks. An additional processing step scores peaks according to an irreproducible discovery rate (IDR), which is a measure of consistency between replicate experiments. Bins are labeled positive if they overlap a peak that meets the IDR threshold of 5%. The IDR scores are not always monotonically associated with the fold-changes. Nevertheless, we expect that performance scores from the fold-change signal classifiers should serve as overly optimistic upper bounds for benchmarking. Commensurate with these expectations, the auPR scores of the FactorNet models are less than, but positively correlative with, the respective auPR scores of the ChIP-seq fold-change signal classifiers (Figure 2A). Interestingly, this pattern does not extend to the auROC scores, and in more than half of the cases the FactorNet auROC scores are greater (Figure 2B). These results are consistent with previous studies that showed the auROC can be unreliable and overly optimistic in an imbalanced class setting (Saito and Rehmsmeier, 2015), which is a common occurrence in genomic applications (Quang and Xie, 2016), motivating the use of alternative measures like the auPR that ignore the overly abundant true negatives.

**Figure 2.**
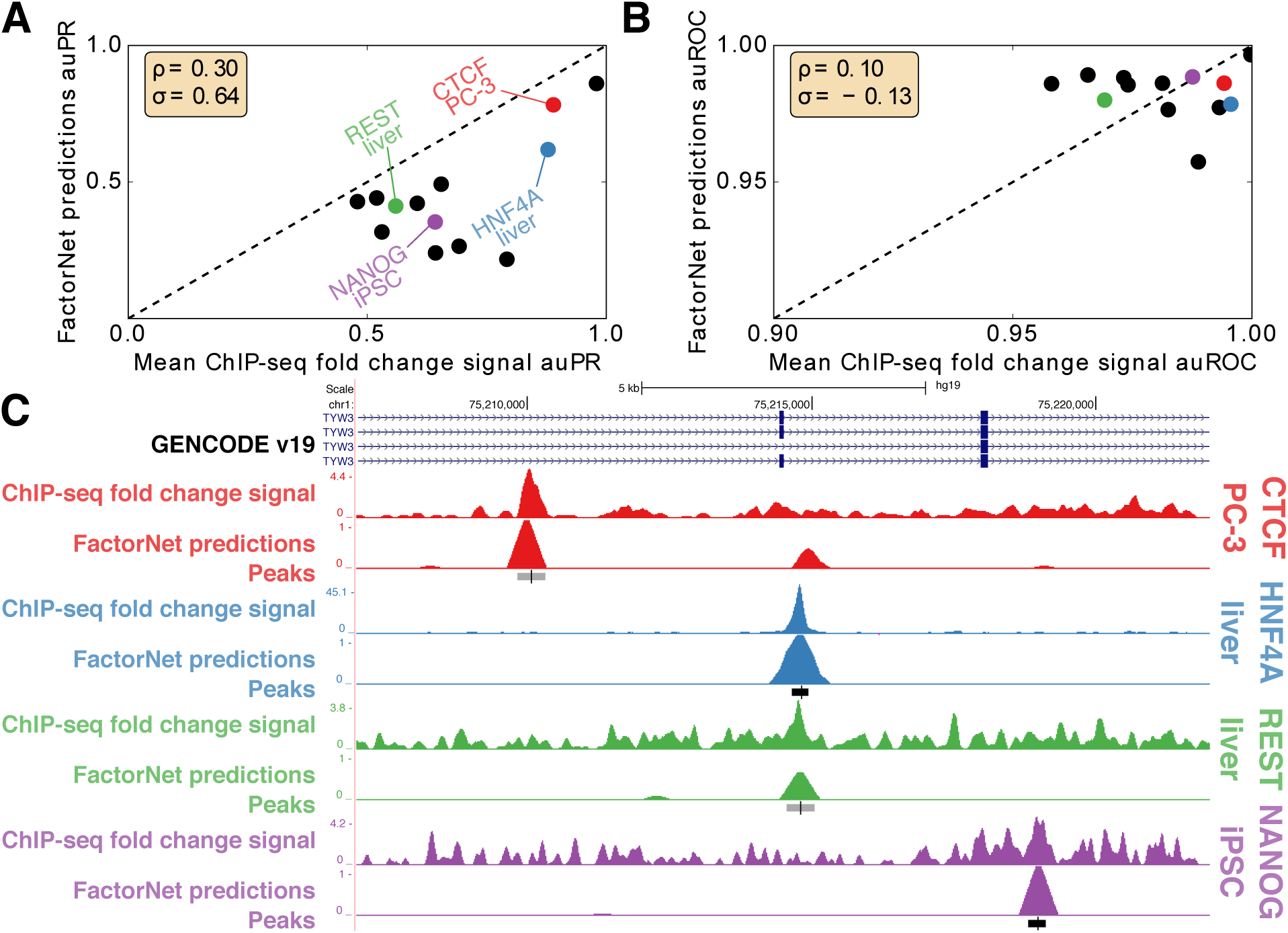
Predictive performance and ChIP-seq signal varies across TF/cell-type pairs. Scat-terplots compare *(A)* auPR and *(B)* auROC scores between FactorNet predictions and mean ChIP-seq fold change signal. Each marker corresponds to one of the 13 final ranking TF/cell type pairs. Spearman (*ρ*) and Pearson (*σ*) correlations are displayed in each plot. *(C)* Genome browser (Kent et al., 2002) screenshot displays the ChIP-seq fold change signal, FactorNet predictions, and peak calls for four TF/cell type pairs in the TYW3 locus. Confidently bound regions are more heavily shaded than ambiguously bound regions.

We can also visualize the FactorNet predictions as genomic signals that can be viewed alongside the ChIP-seq signals and peak calls (Figure 2C and S1). Higher FactorNet prediction values tend to coalesce around called peaks, forming peak-like shapes in the prediction signal that resemble the signal peaks in the original ChIP-seq signal. The visualized signals also demonstrate the differences in signal noise across the ChIP-seq datasets. The NANOG/iPSC ChIP-seq dataset, for example, displays a large amount of signal outside of peak regions, unlike the HNF4A/liver ChIP-seq dataset which has most of its signal focused in peak regions.

The ENCODE-DREAM challenge data, documentation, and results can be found on the Challenge homepage: https://www.synapse.org/ENCODE.

### Interpreting neural network models

Using the same heuristic from DeepBind (Alipanahi et al., 2015) and DanQ (Quang and Xie, 2016), we visualized several kernels from a HepG2 multi-task model as sequence logos by aggregating subsequences that activate the kernels (Figure 3A). The kernels significantly match motifs associated with the target TFs. Furthermore, the aggregated DNase I signals also inform us of the unique “footprint” signatures the models use to identify true binding sites at single-nucleotide resolution. After visualizing and aligning all the kernels, we confirmed that the model learned a variety of motifs (Figure 3B). A minority of kernels display very little sequence specificity while recognizing regions of high chromatin accessibility (Figure 3C).

**Figure 3.**
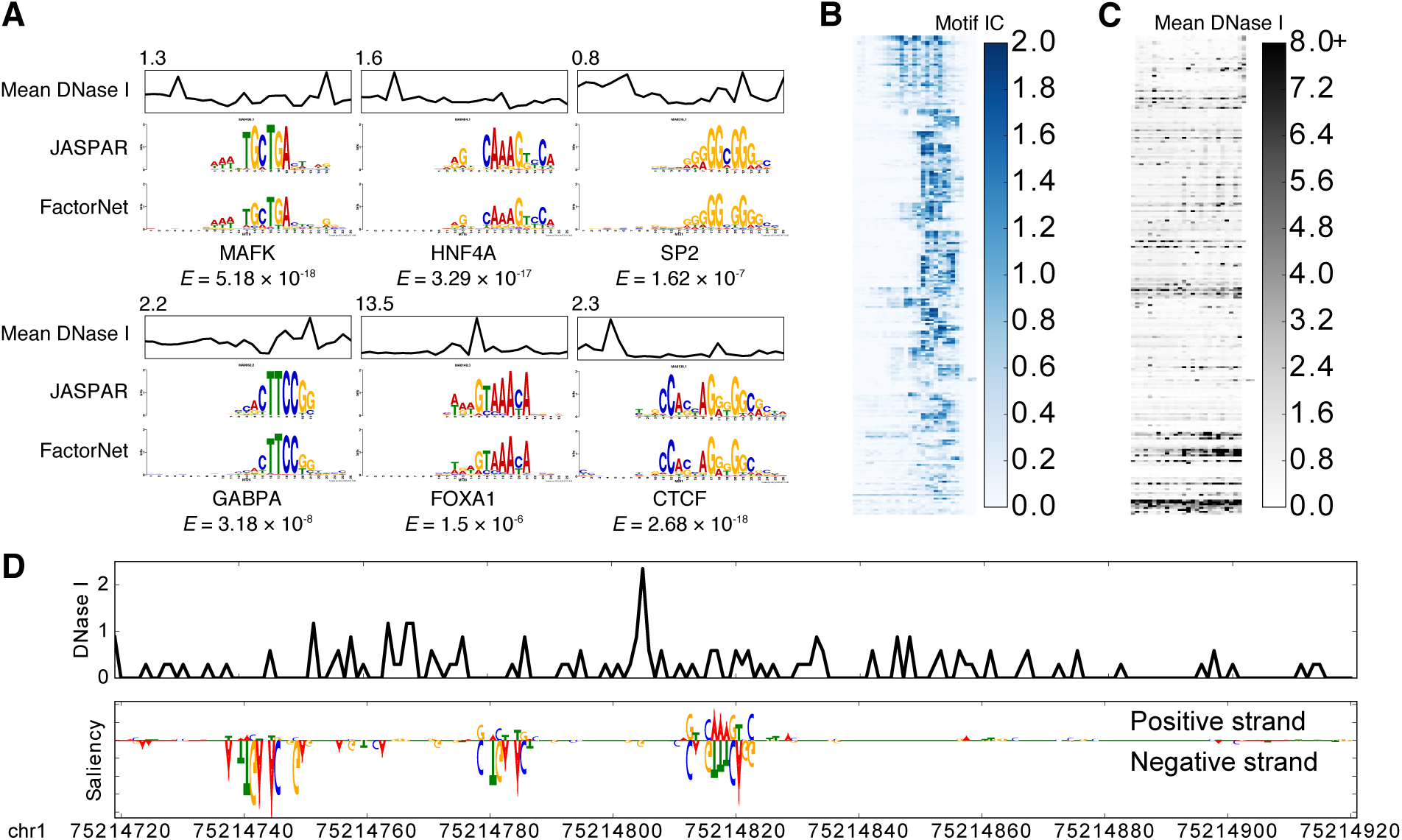
Visually interpreting FactorNet models. *(A)* Network kernels from a HepG2 multi-task FactorNet model are converted to sequence logos and aligned with motifs from JASPAR (Mathelier, A. et al., 2016) using TOMTOM (Gupta et al., 2007). Mean normalized DNase I cleavage signals and their maximum values are displayed above the aligned logos. *E*-values measure similarity between query and target motifs, corrected for multiple hypothesis testing. All kernels are converted to sequence logos and aligned with RSAT (Medina-Rivera et al., 2015). The heatmaps are ordered by this alignment and colored according to the motif information content (IC) *(B)* or mean DNase I cleavage signal *(C)* at each nucleotide position. *(D)* Normalized liver DNase I cleavage signal and saliency maps of aligned stranded sequences centered on the summit of a liver HNF4A peak in the TYW3 locus (Figure 2C). Negative gradients are converted to zeros. We visualized saliency maps with the DeepLIFT visualizer (Shrikumar et al., 2017)

Saliency maps are another common technique of visualizing neural network models (Simonyan et al., 2013). To generate a saliency map, we compute the gradient of the output category with respect to the input sequence. By visualizing the saliency maps of a genomic sequence, we can identify the parts of the sequence the neural network finds most relevant for predicting binding, which we interpret as sites of TF binding at single-nucleotide resolution. Using a liver HNF4A peak sequence and HNF4A predictor model as an example, the saliency map highlights a subsequence overlapping the summit that strongly matches the known canonical HNF4A motif, as well as two putative binding sites upstream of the summit on the reverse complement (Figure 3D).

### Data variation influences predictive performance

In the cases for which two or more testing cell types are available for the same TF, we also observe some rather large disparities in performance. With the exception of FOXA1, FactorNet consistently performs poorer for liver than for other cell types, the difference in auPR reaching as much as 33.5% in the case of EGR1 (Table 1). Variation in data quality across cell type-specific datasets may partially explain these performance differences. The DNase-seq data, which is arguably the most informative cell type-specific feature for binding prediction, widely varies in terms of sequencing depth and signal-to-noise ratio (SNR) across the cell types, which we measure as the fraction of reads that fall into conservative peaks (FRiP) (Figure S2A). Notably, liver displays the lowest SNR with a FRiP score of 0.05, which is consistent with its status as a primary tissue; all other cell types are cultured cell lines.

To further scrutinize the effect data variation has on performance, we trained several FactorNet single-task models and plotted the learning curves to monitor for overfitting (Figure S2B). Learning curves trace the predictive performance of neural networks on training and validation sets. They are useful for identifying signs of overfitting, a common problem in machine learning. These learning curves focus on the GM12878 and HeLa-S3 cell types, using one cell type for training and the other as a validation set. We selected these two cell types because they are the only two reference cell types for E2F1, which FactorNet performed particularly poor on. In addition, the HeLa-S3 DNase-seq data read count and FRiP score are both almost twice that of the read count and FRiP score for the GM12878 DNase-seq data.

From the learning curves of the E2F1 model trained on GM12878, we observe evidence of overfitting. The HeLa-S3 cross-cell type validation loss reaches a minimum value within four training epochs, after which it increases until it reaches a steady state value. In contrast, the GM12878 within-cell type validation loss steadily decreases past the first four epochs and remains much smaller than the HeLa-S3 validation loss throughout training. At first, we speculated the gap to be caused by the differences in the cell type DNase-seq data; however, based on the learning curves for other TFs, this may not necessarily be the sole reason. In the cases of GABPA and TAF1, the differences in validation losses is much smaller. One possible explanation for these results is the differences in the ChIP-seq protocols between the GM12878 and HeLa-S2 datasets. Unlike the other three TFs, the GM12878 and HeLa-S3 E2F1 ChIP-seq datasets were generated using two different antibodies: ENCAB037OHX and ENCAB000AFU, respectively. Both ZNF143 ChIP-seq datasets were generated using the same antibody (ENCAB000AMR), but the model trained on HeLa-S3 displays an unusually high validation loss difference. We speculate this is because the GM12878 ZNF143 ChIP-seq dataset was generated using both single-end 36 bp and paired-end 100 bp reads while the HeLa-S3 ZNF143 ChIP-seq dataset was generated using only single-end 36 bp reads. Given that paired-end 100 bp reads can map to genomic regions that are unmapable for the shorter 36 bp reads, we suspect that differences in read types can introduce significant dataset-specific artifacts.

Given the differences in the GM12878 and HeLa-S3 E2F1 ChiP-seq datasets resulting from the use of different antibodies, we investigated whether a model exclusively trained on one cell type could improve our predictive performance for the K562/E2F1 testing set. To do so, we retrained single-and multi-task models exclusively on either GM12878 or HeLa-S3 and evaluated cross-cell type binding performance on the E2F1/K562 testing set. In contrast, the E2F1 model used at the conclusion of the Challenge was trained on data from both reference cell types. The K562 E2F1 ChIP-seq dataset was generated using the antibodies ENCAB037OHX and ENCAB851KCY, the former of which was also used for GM12878. Hence, we expect that the GM12878 model would be a better predictor for K562 E2F1 binding sites than the other two models, which we find to indeed be the case (Figure S2C-D). Although we managed to improve upon our previous E2F1 model, the cross-cell type performance for E2F1 is still inadequate, especially compared to TFs like CTCF. Predicting binding for TFs in the E2F family is notoriously difficult because members of this protein family share almost identical binding motifs, which in turn makes distinguishing between multiple members of the same family difficult. For TFs that are part of a large family sharing similar sequence binding preference, we conjecture that performance will be limited regardless of the choice of cell type or antibody.

### Comparing single-and multi-task training

Although a thorough comparison between single-and multi-task training is beyond the scope of this paper, our results on E2F1 show that single-and multi-task models can differ in performance. Specifically, the cross-cell type auPR of the single-task GM12878 model is more than 10% greater than that of its respective multi-task model (Figure S2C and Figure S2D). To the best of our knowledge, the cross-cell type performance of each training method depends on the TF/cell type pair. For example, when we retrained single-task and multi-task models for NANOG using H1-hESC as a reference cell type and evaluated the models on iPSC, the multi-task model’s auPR score is over 16% greater than that of the single-task model (Figure S3).

While we initially assumed that the multi-task training confers an advantage by introducing additional information through the multiple labels, at least in the case of NANOG, there are too many conflating variables to immediately conclude this. One of these conflating variables is the differences in training data between single-task and multi-task models. In our current framework, the multi-task training contains significantly more negative bins per training epoch than the single-task training does to balance the positive bins from multiple ChIP-seq datasets. By increasing the ratio of negative to positive samples per epoch for single-task training, we can close the gap between the two training methods in terms of the auPR score, demonstrating that the selection of negative bins affects predictive accuracy. Moreover, the single-task models each use 654,657 weights and require 30 seconds-5 minutes per training epoch whereas the multi-task models each use 5.4 million weights and require 2-3 hours per training epoch, making the former significantly more efficient than the latter. Regardless of whether single-or multi-task training is advantageous, ensembling predictions from both model types can yield significant improvements in performance. It should be noted this pattern did not hold true for the case of training on HeLa-S3 data and evaluating on E2F1/K562 (Figure S2D), and we speculate that the difference in antibodies may explain this discrepancy.

## Discussion

In this work, we introduced FactorNet - an open source package to apply stacked convolutional and recurrent neural networks for predicting TF binding across cell types. While RNNs are computationally expensive to train, especially compared to CNNs, FactorNet incorporates several heuristics to significantly speed up model training and improve predictive performance. Using data from the ENCODE-DREAM Challenge, we demonstrated how our model can effectively integrate cell type-specific data such as DNase-seq to generalize TF binding from reference cell types to testing cell types. As of the conclusion of the Challenge, FactorNet is one of the top performing binding prediction models.

Through our post-Challenge analyses, we gained insights into the variables that affect predictive performance, allowing us to propose strategies for improving the model. First, we observed that the predictive performance widely varied over all TF/cell type pairs, especially in terms of the auPR metric. By leveraging the original ChIP-seq fold change signal, we established upper bounds for the auPR metric for each final ranking TF/cell type pair. These bounds also correlate positively with auPR scores from FactorNet predictions, showing that a large amount of the variation in predictive performance can be attributed to the noise in the original ChIP-seq signal (Figure 2A). We expect that predictive performance for many TF/cell type pairs can be improved by redoing experiments with higher quality antibodies. Alternatively, ChIP-exo, a modification of ChIP-seq that uses exonucleases to degrade contaminating non-protein-bound DNA fragments (Rhee and Pugh, 2011), may improve the quality of ChIP signals. Next, we investigated the variation in the DNase-seq datasets. We found that the DNase-seq datasets greatly differ in terms of sequencing depth and SNR (Figure S3). While we do correct for the variation in sequencing depth by normalizing the cleavage signals to 1x coverage, we do not correct for the variation in the SNR. The performance lost is most staggering for the liver cell type, which has the DNase-seq dataset with the lowest SNR. However, differences in DNase-seq SNR do not fully account for differences in predictive performance. By studying several within-and cross-cell type validation curves, we also concluded that differences in antibodies and read lengths can introduce significant dataset-specific biases (Figure S2B). Accordingly, we can improve performance by omitting less compatible cell type datasets (Figure S2C-D).

We also compared single-and multi-task training frameworks. Several deep learning methods, including DeepSEA and Basset, primarily use multi-task training, which involves assigning multiple labels, corresponding to different chromatin markers, to the same DNA sequence. The authors of these methods propose that the multi-task training improves efficiency and performance. FactorNet supports both types of training. Our results do not entirely favor one method over the other. For the K562/E2F1 cross-cell type testing set, the GM12878 single-task model outperformed GM12878 multi-task model (Figure S2C); however, for the NANOG/iPSC cross-cell type testing set, the H1-hESC multi-task model outperformed the H1-hESC single-task model (Figure S3). In the latter case, the performance gap can be narrowed by changing the proportion of negative to positive training samples in the single-task framework, suggesting that any additional gain granted by the multiple labels is eclipsed by the choice of negative sets. Nevertheless, ensembling single-and multi-task models together appears to be an effective method of improving predictive performance, at least if antibodies and read lengths are kept consistent.

Another avenue we can explore for improving the model is hyperparameter tuning. We selected the hyperparameters for the models in this work arbitrarily for demonstration and uniformity purposes (Table S1-S3). Although we have not yet implemented them, distributed computing hyperparameter tuning algorithms (Bergstra et al., 2013) can systematize hyperparameter selection and improve performance.

One of the chief criticisms of neural networks is that they are “black box” models. While neural networks can achieve great performances in predictive tasks, the exact reasons for why this is the case is not always entirely clear. In contrast to these criticisms, we can visualize and interpret aspects of the FactorNet model. By converting network kernels to motifs, we show that FactorNet can recover motifs that are known to contribute to binding (Figure 3A). DNase I footprint patterns help discriminate true binding sites from putative sites that simply match a motif. Previous TF binding prediction methods, such as Centipede, require users to supply motifs. FactorNet relaxes this strong assumption and essentially performs *de novo* motif discovery during the learning process to identify the sequence patterns that are most useful for binding prediction. Saliency maps can also help elucidate the complex regulatory grammar that govern TF binding by visualizing the spatial positions and orientations of multiple binding sites that work together to recruit TFs (Figure 3D).

Our adherence to standardized file formats also makes FactorNet robust. For example, FactorNet can readily accept other genomic signals that were not included as part of the Challenge but are likely relevant to TF binding prediction, such as conservation and methylation. Along these same lines, if we were to refine our pre-processing strategies for the DNase-seq data, we can easily incorporate these improved features into our model as long as the data are available as bigWig files (Kent et al., 2010). Other sources of open chromatin information, such as ATAC-seq (Buenrostro et al., 2015) and FAIRE-seq (Giresi et al., 2007), can also be used to replace or complement the existing DNase-seq data. In addition, FactorNet is not necessarily limited to only TF binding predictions. If desired, users can provide the BED files of positive intervals to train predictive models for other markers, such as histone modifications. As more epigenetic datasets are constantly added to data repositories, FactorNet is already in a prime position to integrate both new and existing datasets.

In conclusion, FactorNet is a very flexible framework that lends itself to a variety of future research avenues. The techniques that we introduced in this paper will also be useful for the field of machine learning, especially since neural network models are becoming increasingly popular in genomics. Some of the design elements of FactorNet were motivated by the specific properties inherent in the structure of the data. Many of these properties are shared in data found in other applications of machine learning. For example, the directional nature and modularity of DNA sequences prompted us to search for a model that can discover local patterns and long-range interactions in sequences, which led us to ultimately select a hybrid neural network architecture that includes convolution and bidirectional recurrence. Natural language processing problems, such as topic modeling and sentiment analysis, can also benefit from such an architecture since language grammar is directional and modular. Another unique aspect of the data that guided our design is the double strandedness of DNA, which prompted us to adopt a Siamese architecture to handle pairs of input sequences (Figure 1). Protein-protein interaction prediction also involves sequence pairs and would likely benefit from a similar framework. Our heuristics for reducing training time and computational overhead will serve as useful guidelines for other applications involving large imbalanced data, especially if recurrent models are utilized. We therefore expect that FactorNet will be of value to a wide variety of fields.

## Methods

### ENCODE-DREAM Challenge dataset

The ENCODE-DREAM Challenge dataset is comprised of DNase-seq, ChIP-seq, and RNA-seq data from the ENCODE project or The Roadmap Epigenomics Project covering 14 cell types and 32 TFs. All annotations and pre-processing are based on hg19/GRCh37 release version of the human genome and GENCODE release 19 (Harrow et al., 2012). Data are restricted to chromosomes X and 1-22. Chromosomes 1, 8 and 21 are set aside exclusively for evaluation purposes and binding data were completely absent for these three chromosomes during the Challenge. TF binding labels are provided at a 200 bp resolution. Specifically, the genome is segmented into 200 bp bins sliding every 50 bp. Each bin is labeled as bound (B), unbound (U) or ambiguously bound (A) depending on the majority label of all nucleotides in the bin. Ambiguous bins overlap peaks that fail to pass the IDR threshold of 5% and are excluded from evaluation. A more complete description of the dataset, including pre-processing details such as peak calling, can be found in the ENCODE-DREAM Challenge website (https://www.synapse.org/ENCODE).

### Evaluation

The TF binding prediction problem is evaluated as a two-class binary classification task. For each test TF/cell type pair, the following performance measures are computed:

1. **auROC**. The area under the receiver operating characteristic curve is a common metric for evaluating classification models. It is equal to the probability that a classifier will rank a randomly chosen positive instance higher than a randomly chosen negative one.
2. **auPR**. The area under the precision-recall curve is more appropriate in the scenario of few relevant items, as is the case with TF binding prediction (Quang and Xie, 2016). Unlike the auROC metric, the auPR metric does not take into account the number of true negatives called.
3. **Recall at fixed FDR**. The recall at a fixed false discovery rate (FDR) represents a point on the precision-recall curve. Like the auPR metric, this metric is appropriate in the scenario of few relevant items. This metric is often used in applications such as fraud detection in which the goal may be to maximize the recall of true fraudsters while tolerating a given fraction of customers to falsely identify as fraudsters. The ENCODE-DREAM Challenge computes this metric for several FDR values.

As illustrated in Figure 1, the FactorNet Siamese architecture operates on both the forward and reverse complement sequences to ensure that both strands return the same outputs during both training and prediction. Although a TF might only physically bind to one strand, this information cannot usually be inferred directly from the peak data. Thus, the same set of labels are assigned to both strands in the evaluation step.

### Features and data pre – processing

FactorNet works directly with standard genomic file formats and requires relatively little pre-processing. BED files provide the locations of reference TF binding sites and bigWig files (Kent et al., 2010) provide dense, continuous data at single-nucleotide resolution. bigWig values are included as extra rows that are appended to the four-row one hot input DNA binary matrix. FactorNet can accept an arbitrary number of bigWig files as input features, and we found the following signals to be highly informative for prediction:

1. **DNase I cleavage**. For each cell type, reads from all DNase-seq replicates were trimmed down to first nucleotide on the 5’ end, pooled and normalized to 1x coverage using deepTools (Ramírez et al., 2014).
2. **35 bp mapability uniqueness**. This track quantifies the uniqueness of a 35 bp subsequence on the positive strand starting at a particular base, which is important for distinguishing where in the genome DNase I cuts can be detected. Scores are between 0 and 1, with 1 representing a completely unique sequence and 0 representing a sequence that occurs more than 4 times in the genome. Otherwise, scores between 0 and 1 indicate the inverse of the number of occurrences of that subsequence in the genome. It is available from the UCSC genome browser under the table wgEncodeDukeMapabilityUniqueness35bp.

In addition to sequential features, FactorNet also accepts non-sequential metadata features. At the cell type level, we applied principal component analysis to the inverse hyperbolic sine transformed gene expression levels and extracted the top 8 principal components. Gene expression levels are measured as the average of the fragments per kilobase per million for each gene transcript. At the bin level, we included Boolean features that indicate whether gene annotations (coding sequence, intron, 5’ untranslated region, 3’ untranslated region, and promoter) and CpG islands (Gardiner-Garden and Frommer, 1987) overlap a given bin. We define a promoter to be the region up to 300 bps upstream and 100 bps downstream from any transcription start site. To incorporate these metadata features as inputs to the model, we append the values to the dense layer of the neural network and insert another dense layer containing the same number of ReLU neurons between the new merged layer and the sigmoid layer (Figure 1).

### Training

Our implementation is written in Python, utilizing the Keras 1.2.2 library (Chollet et al., 2015) with the Theano 0.9.0 (Bastien et al., 2012; Bergstra et al., 2010) backend. We used an NVIDIA Titan X Pascal GPU for training.

FactorNet supports single-and multi-task training. Both types of neural network models are trained using the Adam algorithm (Kingma and Ba, 2014) with a minibatch size of 100 to minimize the mean multi-task binary cross entropy loss function on the training set. We also include dropout (Srivastava et al., 2014) to reduce overfitting. One or more chromosomes are set aside as a validation set. Validation loss is evaluated at the end of each training epoch and the best model weights according to the validation loss are saved. Training sequences of constant length centered on each bin are efficiently streamed from the hard drive in parallel to the model training. Random spatial translations are applied in the streaming step as a form of data augmentation. Each epoch, an equal number of positive and negative bins are randomly sampled and streamed for training, but this ratio is an adjustable hyperparameter (see Table S1 for a detailed explanation of all hyperparameters). In the case of multi-task training, a bin is considered positive if it is confidently bound to at least one TF. Bins that overlap a blacklisted region (ENCODE Project Consortium, 2012) are automatically labeled negative and excluded from training.

### Single-task training

Single-task training leverages data from multiple cell types by treating bins from all cell types as individually and identically distributed (i.i.d.) records. To make single-task training run efficiently, one bin is allotted per positive peak and these positive bins are included at most once per epoch for training. Each epoch, negative bins are also drawn randomly without replacement from the training chromosomes. For example, if we were to train on a single cell type that has 10,000 peaks for a particular TF, then we may train on 10,000 positive bins and 10,000 negative bins each epoch. Ambiguously bound bins are excluded from training.

### Multi-task training

FactorNet can only perform multi-task training when training on data from a single cell type due to the variation of available binding data for the cell types. For example, the ENCODE-DREAM Challenge provides reference binding data for 15 TFs for GM12878 and 16 TFs for HeLa-S3, but only 8 TFs are shared between the two cell types. Unlike the single-task models, which ignore ambiguous bins during training, the multi-task models assign negative labels to the ambiguous bins because of the frequent overlap of confidently and ambiguously bound regions. Compared to single-task training, multi-task training takes considerably longer to complete due to the larger number of positive bins. At the start of training, positive bins are identified by first segmenting the genome into 200 bins sliding every 50 bp and discarding all bins that fail to overlap at least one confidently bound TF site. Each epoch, negative bins are drawn randomly with replacement from the training chromosomes.

### Bagging

Ensembling is a common strategy for improving classification performance. At the time of the Challenge, we implemented a simple ensembling strategy commonly called “bagging submissions”, which involves averaging predictions from two or more models. Instead of averaging prediction probabilities directly, we first convert the scores to ranks, and then average these ranks. Rank averaging is more appropriate than direct averaging if predictors are not evenly calibrated between 0 and 1, which is often the case with the FactorNet models.

### Software availability

Source code is available at the github repository http://github.com/uci-cbcl/FactorNet. In addition to the source code, the github repository contains all models and data used for the ENCODE-DREAM Challenge.

## Acknowledgments

We thank the ENCODE-DREAM challenge organizers for providing the opportunity to test and improve our method. We also thank David Knowles for helping with generating gene expression metadata features.

This work was supported by the National Institute of Biomedical Imaging and Bioengineering, National Research Service Award (EB009418) from the University of California, Irvine, Center for Complex Biological Systems and the National Science Foundation Graduate Research Fellowship under Grant No.(DGE-1321846). Any opinion, findings, and conclusions or recommendations expressed in this material are those of the authors and do not necessarily reflect the views of the National Science Foundation.

## Supporting Information

**Table S1.** Summary and description of the hyperparameters used for the single-task models in Figure S2B.

| Hyperparameter | Value | Description |
| --- | --- | --- |
| -v validchroms | chr3 chr5 chr7 chr10 chr12 chr14 chr16 chr18 chr20 chrX | Sequences on these chromosomes are set aside for validation. |
| -e epochs | 200 (ZNF143, TAF1), 300 (E2F1, GABPA) | Max number of epochs to train before training ends. |
| -ep patience | 200 (ZNF143, TAF1), 300 (E2F1, GABPA) | Number of epochs with no improvement in the validation loss. |
| -lr learningrate | 0.00001 | Learning rate for the Adam optimizer. We decreased it from the default value of 0.001 to smooth the learning curves. |
| -n negatives | 1 | Number of negative bins to sample per positive bin per epoch. |
| -L seqlen | 1000 | Length, in bps, of input sequences to the model. |
| -w motifwidth | 26 | Width, in bps, of the convolutional kernels. |
| -k kernels | 32 | Number of kernels/motifs in the model. |
| -r recurrent | 32 | Number of recurrent units (in one direction) in the model. |
| -d dense | 128 | Number of units in the dense layer in the model. |
| -p dropout | 0.5 | Dropout rate between the recurrent and dense layers. Also the dropout rate between the dense and sigmoid layers. |
| -m metaflag | False | Flag for including cell type-specific meta-data features (usually gene expression). |
| -g gencodeflag | True | Flag for including CpG island and gene annotations. |
| -mo motifflag | True | Flag for initializing two of the kernels to the PWM of the canonical motif (forward and reverse complement). |
| -s randomseed | Varies | Random seed for reproducibility. |

**Table S2.** Hyperparameters used for the multi-task models in Figures 3 and S2-S3. Unspecified values should be assumed to be the same as those found in Table S1.

| Hyperparameter | Value | Notes |
| --- | --- | --- |
| -v validchroms | chr11 |  |
| -e epochs | 20 | Fewer epochs needed for multi-task training due to the large number of training bins. |
| -ep patience | 20 |  |
| -lr learningrate | 0.001 | Default value of 0.001 is sufficient for most applications. |
| -n negatives | 1 |  |
| -g gencodeflag | False | Multi-task training does not currently incorporate any metadata features. |
| -mo motifflag | False |  |

**Table S3.** Hyperparameters used for the single-task models in Figures S2C-D and S3. Unspecified values should be assumed to be the same as those found in Table S1.

| Hyperparameter | Value | Notes |
| --- | --- | --- |
| -v validchroms | chr11 |  |
| -e epochs | 100 | Need more epochs than multi-task training due to fewer positive bins. |
| -ep patience | 20 |  |
| -lr learningrate | 0.001 | Default value of 0.001 is sufficient for most applications. |
| -n negatives | Varies (but usually 1) | In some cases, increasing this value from 1 improves cross-cell type auPR scores for single-task models. |

**Figure S1.**
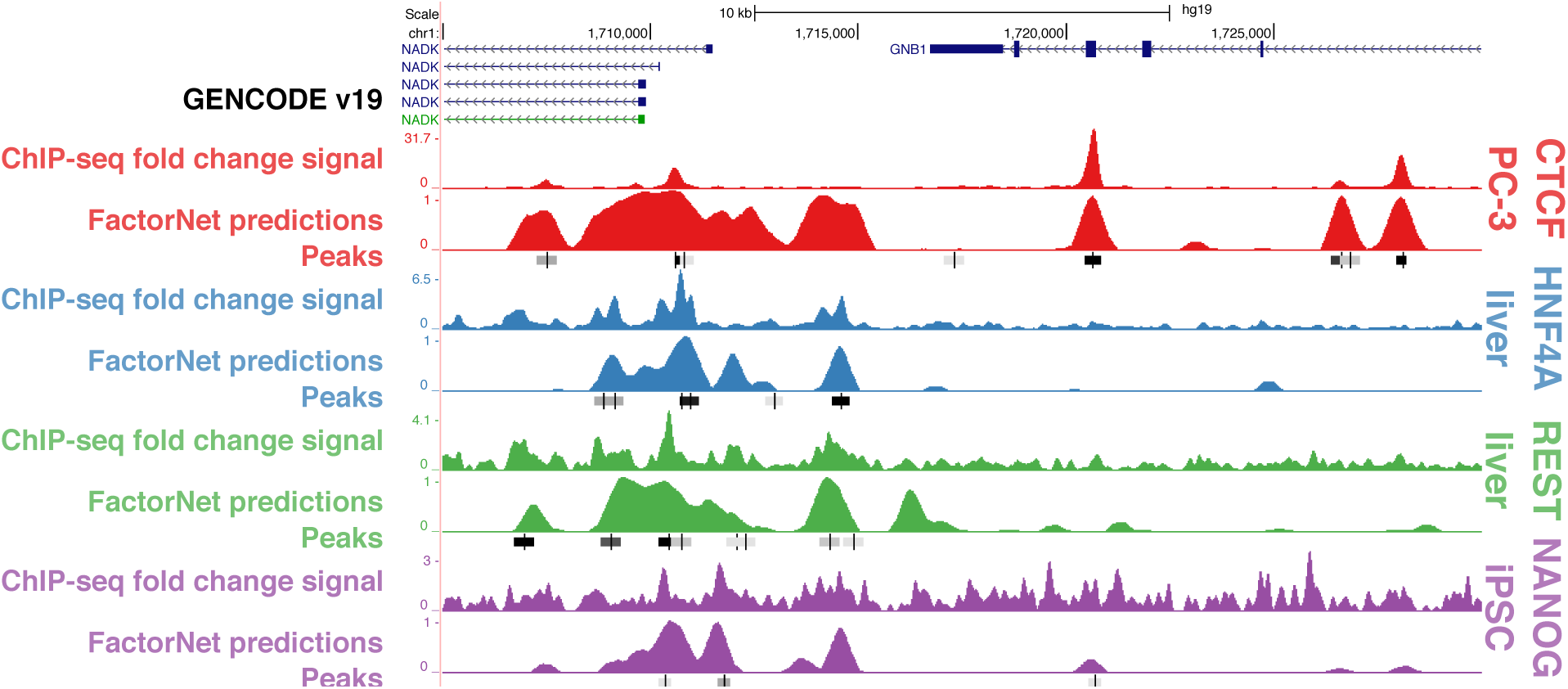
FactorNet cross-cell type predictions are comparable to ChIP-seq signals and peaks. A genome browser shot similar to Figure 2A focusing on the NADK/GNB1 locus.

**Figure S2.**
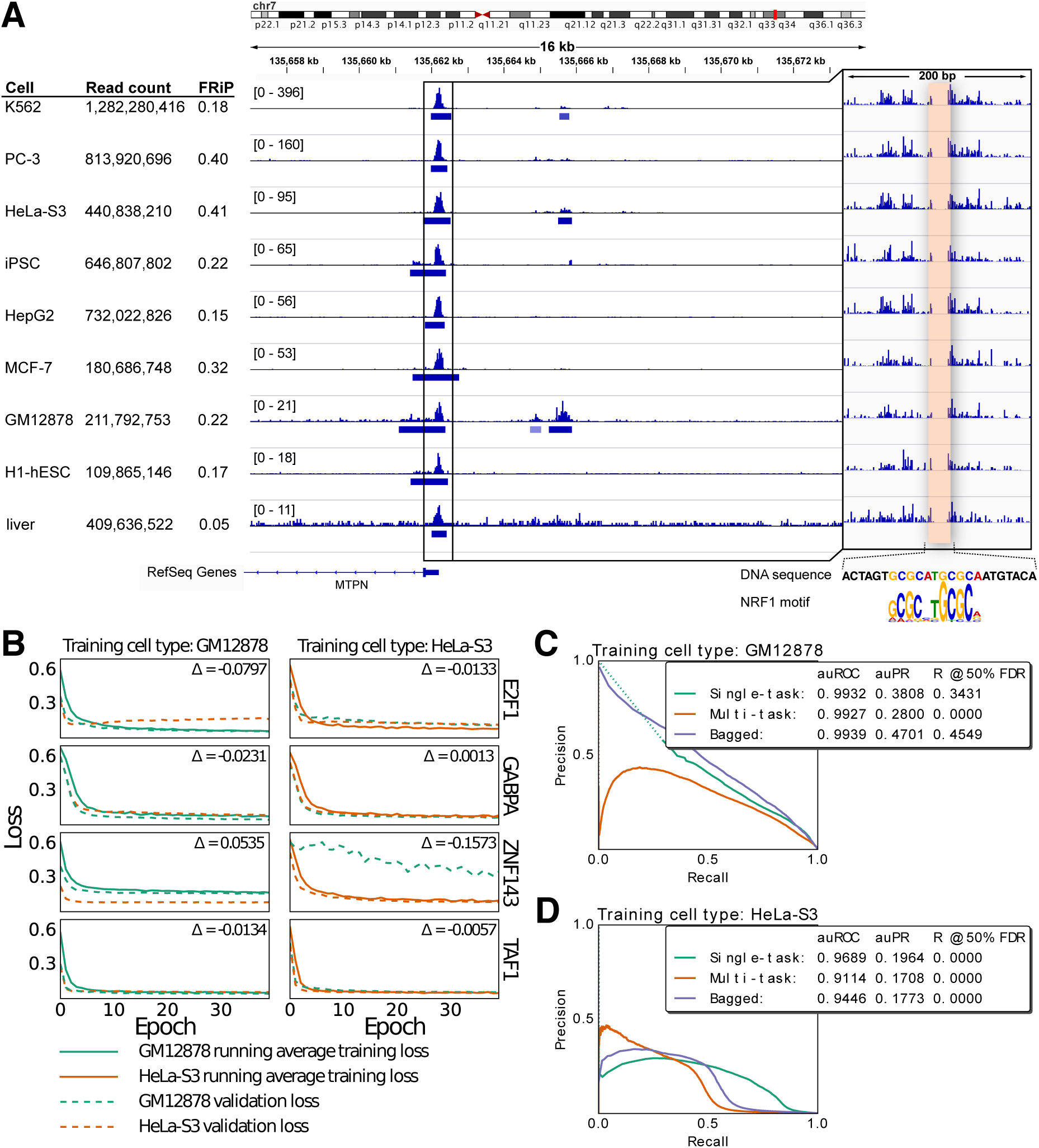
Variation in cell type-specific datasets influence cross-cell type predictive performance. *(A)* IGV (Thorvaldsdóttir et al., 2013) browser screenshot displays pooled DNase I cleavage signal and conservative DNase-seq peaks for eight cell types. The inset is a magnified view at the MTPN promoter, a known NRF1 binding site. *(B)* Each plot displays learning curves of single-task models trained on either GM12878 or HeLa-S3. We generated within-and cross-cell type validation sets by extracting an equal number of positive and negative bins from the validation chromosomes. The difference between the smallest within-and cross-cell type validation losses are displayed in each plot. *(C and D)* Precision-recall curves of single-and multi-task models evaluated on the E2F1/K562 testing set trained exclusively on either GM12878 or HeLa-s3. Dotted lines indicate points of discontinuity. Model weights were selected based on the within-cell type validation loss on chr11. We generated single-task scores by bagging scores from two single-task models initialized differently. Final bagged models ensemble respective single-and multi-task models.

**Figure S3.**
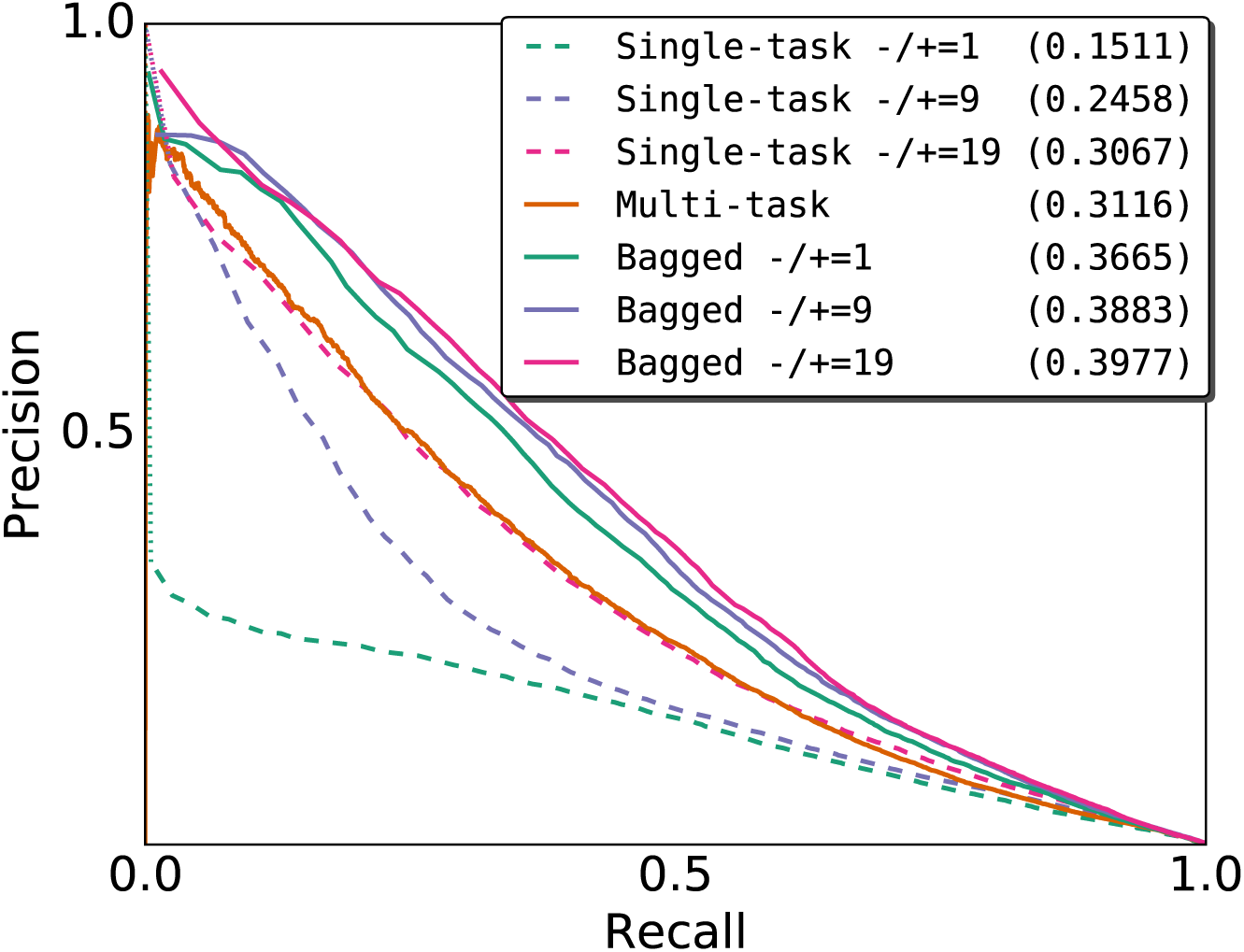
Comparison of single-and multi-task training. Cross-cell type precision-recall curves of single-task and multi-task NANOG binding prediction models trained on H1-hESC and evaluated on iPSC. Model weights were selected based on the within-cell type validation loss on chr11. We generated single-task scores by bagging scores from two single-task models initialized differently. The three single-task models differ in the ratio of negative-to-positive bins per training epoch. The bagged models are the rank average scores from the multi-task model and one of the three single-task models. auPR scores are in parentheses. Both training and testing ChiP-seq datasets use the ENCAB000AIX antibody.

